# Structure Specific Neuro-toxicity of α-Synuclein Oligomer

**DOI:** 10.1101/2020.10.11.334979

**Authors:** Animesh Mondal, Sandip Dolui, Sukhamoy Dhabal, Ashish Bhattacharjee, Nakul C Maiti

## Abstract

Parkinson’s disease (PD) is linked to α-synuclein (aS) aggregation and deposition of amyloid in the substantia nigra region of the brain tissues. Recent reports suggested that oligomeric assembly structure could be neurotoxic to neuronal cells. In the current investigation we produced two distinct classes of aS oligomers and link the protein conformation state and stability to neuronal cell toxicity. Natural oligomers (NO) enriched with alpha-helical folds are produced in storage of aS at below −20°C for 7 days. Induced oligomer (IO), often observed in the aggregation pathway of aS were made incubating the protein solution at 37°C. Natural oligomers remained stable and did not transform into β-sheet rich amyloid fiber and exhibited higher toxicity (80% cell death) compared to induced oligomers. Natural oligomers were ovular shape and the size ranged between 4-5.5 nm. It maintained significant number (∼ 60%) of residues in α-helical conformational space. However, initiation of hydrophobic zipping with beta sheet conformation was evidenced in induced oligomer (IO) and a lesser number residues (45%) remained with preference to α-helical secondary structure. Hydrophobic collapse leads the transformation of IO into thermodynamically most stable β-sheet rich amyloid fibril. Molten globule like secondary structure stabilized by H-bonding in natural oligomers caused enhanced stability and cellular toxicity compared to induced oligomer. Thus off-pathway/natural oligomers could be plausible reason of neuronal cell death and possible cause of Parkinson’s disease.

## Introduction

Parkinson’s disease (PD) is a neurodegenerative disorder and largely characterized by progressive movement disorder with symptoms of tremor, rigidity of limbs and impaired balance.^1^ These bedevilled etiology are arises due to the death of dopamine producing neuron-cells in the substantia nigra region of the brain. In PD and similar other neurodegenerative disorders a key finding is the engulfment of amyloid inclusions, known as ‘lewy bodies’ or ‘lewyneurities’ enriched with protein aggregates. The main component of these ‘lewy bodies’ is α-synuclein (aS) along with other proteins such as p62 and ubiquitin.^2^ However, from different investigation it is established that aS is the key component in this inclusion bodies of patients suffering from PD. Several missense mutation as such as A53T, E46K, A30P are also found to be involved in the pathogenic amyloids found in PD.^3^ However, the mechanistic details behind the cause are still a matter of dilemma.

aS is a member of the synuclein family that also includes β and γ-synuclein often found to localize at the nuclear envelope and at presynaptic nerve terminals.^4^ However the real function or biological role of the protein is poorly understood. It has been suggested that aS may have role in synaptic neuro-transmission, and may regulate synaptic membrane biogenesis.^5^ However, the native structure of the protein *in-vivo* is illusive and may remain associated with lipid molecules preferably with α-helical structure.^6–8^

Although, the protein *in vitro* solution condition is highly fluctuating. It is highly dynamic and capable of adopting different conformation depending upon its surrounding environments. The solution state NMR and Raman spectroscopic analysis firmly established its disordered state without having a compact and stable globular fold.^3,9–12^ In addition, under appropriate solution condition the protein transformed into β-sheet enriched fibrillar structure. During the formation of fibrillar aggregates of aS, several *in-situ* measurements observed various intermediate species such as short-lived globular like oligomer, annular oligomers, diffusible protofibrils.^13–16^ While the understanding of amyloid pathogenesis is highly debated, these early formed oligomeric structures are believed to be nurotoxic.^17^ Several studies observed that early formed oligomeric assembly structures are toxic to model systems. The question arises how these oligomeric species could be neurotoxic and associated with the pathology of amyloid diseases. It is suggested that with spherical shape or pore-like structures, oligomers/protofibrils may disrupt the cell membrane very easily.^18^ Also it is very unclear how these intermediate and their morphology entities are transformed to mature fibrils and, an important another argument is whether misfolding into β-rich structures precedes or follows aggregation.^19,20^

In solution monomeric aS exists as a random conformation and upon binding with the membrane it can attain helical^21^ conformation as a apolipoprotein it contains repetitive motif of KTKEGV. Disorder-to-order transitions may have a crucial role to play in macromolecular recognition. There are numerous examples of protein-protein, protein-nucleic acid, and protein-ligand interactions involving disordered protein segments. It has been postulated that disorder-to-order transitions provide a mechanism for uncoupling binding affinity and specificity, thereby permitting weak but highly specific interactions, or conversely, strong but relatively nonspecific interactions. Such monomeric form interacts among themselves and form oligomers which is a transient species. These oligomeric species are capable of disrupting the membrane integrity^22,23^ such oligomers exhibit distinct levels of toxicity in neuronal cells. Perturbing the structural elevation of aS during inclusion body formation requires precise identification of the species that are highly toxic and capable of promoting the formation of newer aggregates. Several studies indicate there is a variation in the degree of pathogenicity imparted by different species of oligomers and fibers^24,25^.

In animal model the potential role of aS fibrill causing neuronal cell damage was illustrated^26–28^. However, very few reports are available regarding the structural elucidation of oligomeric species. These oligomeric species being a transient or intermediate species can cause high damage to the cell while transmitting from cell to cell during disease progression.^29–31^ Such oligomers exert their cytotoxic effects by creating mitochondrial damage, ER stress and activating proinflammatory cascade. When oligomers create pore like structure it results in the alternation of calcium release and augment neurodegeneration.^32,33^ Several studies suggests that soluble oligomers are more toxic compared to the insoluble aggregates that are easily identifiable with light microscope^34^. Hitherto, very little is known about the structure of these toxic species and their correlation with structural propensity and toxic behaviour. The transient nature of the on pathway oligomeric species converts it rapidly into insoluble amyloid fibrils. From several reports striking morphologies of the aS oligomeric species is well versed; such as spheres, ring and non-fibrillar fillaments^35,36^. Although the morphology and characteristic features of these oligomeic species can be altered in the presence of different cellular microenvironment. In recent report it has been shown that in presence of metal ion such as Iron promotes antiparallel oligomerization^37^. Interestingly, the addition of some metals, lipids and chaperons to monomeric as well as fibrillar form of protein can usher the formation of intermediate species which are characterised as stable in its form and disorder in its conformation. Experimental studies indicate that conformational changes of aS can decree pathology and crucial in disease progression.^38^Treatments of PD by targeting fibrills causes more toxicity to the cell and for that reason characterization of the toxic oligomeric species could probe new dimension in therapeutics.

This study characterized two different forms of oligomeric species by extrapolating Raman spectroscopy. This technique enjoys many important advantages while characterising bond vibration and secondary structural propensity. The feasibility of this method is not hindered by the higher molecular assembly or size of the molecules. Raman spectroscopic characterization can be applied to protein in various states; aqueous solutions, precipitated fibrils, amorphous aggregates, solids and crystals. Raman studies illustrated that in a native condition a fraction of extended structure exists as poly proline-II in native condition^39^. The pliability of the Raman spectroscopy allowed us to gather spectroscopic signals of the two different form of oligomeric species using amide-I and amide-III vibration region along with the full fingerprint region of the spectrum.

It permits to measure the intrinsic molecular vibration of the amide region of the without any external probe. Besides the characterization of the amide-I and other amide bands, there are additional features in Raman spectra that permit the environment of other amino acids side chains to be characterized in their different physical states. Prior Raman studies on aS is very limited. Secondary structural conformations of the oligomeric species were evaluated in a non-destructive method and finally correlated its structural propensity and toxic form in neuronal cell line. Our study is well validated with other biophysical techniques such as CD, AFM and fluorescence study. The study report detailed characterization of the toxic oligomeric species of which one form is associated with amyloid fibrillation and other without forming amyloid fibrills capable of exerting its highly neurotoxic propensity. Lansbury et.al suggests that spheroidal oligomers of aS are to be rich in β-sheet structure^40^. However, we observed some higher amount of helicity in one of the oligomeric species and the other is rich in β-sheet conformation as depicted by earlier studies. The results of this study bestow the basis for more holistic understanding of the ensemble of the oligomeric species and its behavioural peculiarity towards nerve cell and tissues.

## Results and Discussions

In the current investigation the study focused on the structural intricacy of α-synuclein (aS) oligomers and their correlation with neurotoxicity. Fresh aS solution was made using recombinant technology. aS was expressed and purified using established protocols and then its purity was judged in by running SDS PAGE and MALDI as showed in upper panel of the Figure S1b. For making natural oligomer (NO) the monomeric fraction of the purified protein was concentrated and lyophilized. Again it was re-suspended in buffer and subsequently it was passed through gel filtration column. From the solution monomer and natural oligomers were isolated at particular fraction of time (Figure S1a). During elution natural oligomers were eluted first followed by monomeric portion through the size exclusion column. The on pathway oligomers or induced oligomers (IO) were made by incubating the monomeric protein solution (200 µM) at 37°C under constant shaking (750 rpm). Details are given in the materials and the method section.

To determine the stability of the two types of oligomeric species (NO and IO) SDS-PAGE and native PAGE were performed. Interestingly, both of the oligomers degraded when run through SDS-PAGE and converted to monomeric form of aS (Figure S1c, left panel). This indicated both of them were not SDS resistant and very much prone to degradation. When performed native gel electrophoresis of the two oligomeric species both of them remained in the stacking portion of the gel (Figure S1c right panel). However, a consistent observation was that the induced oligomer formed smears during migration but it failed to migrate nicely as monomeric form of the protein (Figure S1c, right panel). Natural oligomers did not form smear and remained in the stacking portion of gel as a stable form. It suggested better stability and uniform population compared to the induced oligomers.

The surface morphological features of the two type oligomers were further investigated by atomic force microscopy (AFM). Figure 1 displays AFM image of induced aS oligomers. The oligomer (IO) was made from aS incubating at 37°C under constant shaking (750 rpm) for 2 days. It was drop casted on freshly cleaved mica by removing solution from the incubated aS solution at a particular time interval. The induced oligomers (IO) were found to be spheroidal in nature. The height of the spheroidal oligomers found to be 1-2 nm and its diameter was between 25-35 nm. However, after prolonged incubation of 4 days these oligomers gradually changed and converted itself to higher order protofibril/fibrillar aggreagtes. The height of this fibrillar assembly was ∼5 nm and its length varied up to few micrometers. A dense network of fibrills were noticed which served as a precursor of amyloid aggregates. This observation denoted that induced oligomeric species were on pathway assembly structure that eventually transformed into mature fibrils. As depicted in our results suggested that early aggregation of aS progress through molecular assembly of monomer to oligomer, and subsequent interactions among themselves hierarchically generate higher order fibrillar structure (Scheme 1).^41^ It has been observed that amyloid fibrills can be formed from a wide range of proteins like aS, however, the common feature prevailing within this amyloid fibrills is the core architecture of cross β structures. aS first polymerize among themselves as oligomers and these oligomers further proceeds through their course of aggregation to form protofibrills. The fibrills typically results from the assembly of protofilaments/protofibrils which wrap around each other and generates a twisted structure. Information from numerous studies indicates that β strands align perpendicular to each other along the fibrillar axis. However, from several deposited molecular structure it is well established that both intramolecular and intermolecular hydrogen bonds play pivotal role in interaction of β sheets^42^. However, in the current investigation the major goal was to precisely determine the surface morphology and the structural component and to correlate them to the neuronal toxicity caused by the oligomers. Further aim was to define the structure and conformation of natural oligomers which failed to produce fibrillar structure (as discussed below) and compared the toxic effect with the IO. Natural oligomer (NO) showed slightly different surface morphology (Figure 1 left panel). Shape of the oligomers appeared to be ovular and its diameter varied from 20-25 nm which was slightly smaller than induced oligomers. Freshly prepared oligomeric population possessed cross sectional height of 4-5.5nm; However, incubation of these oligomers for 4 days under previous condition; the oligomers became slightly bigger in size. Interestingly, it did not attain any fibrillar morphology even after incubating for four days.

**Figure 1:**
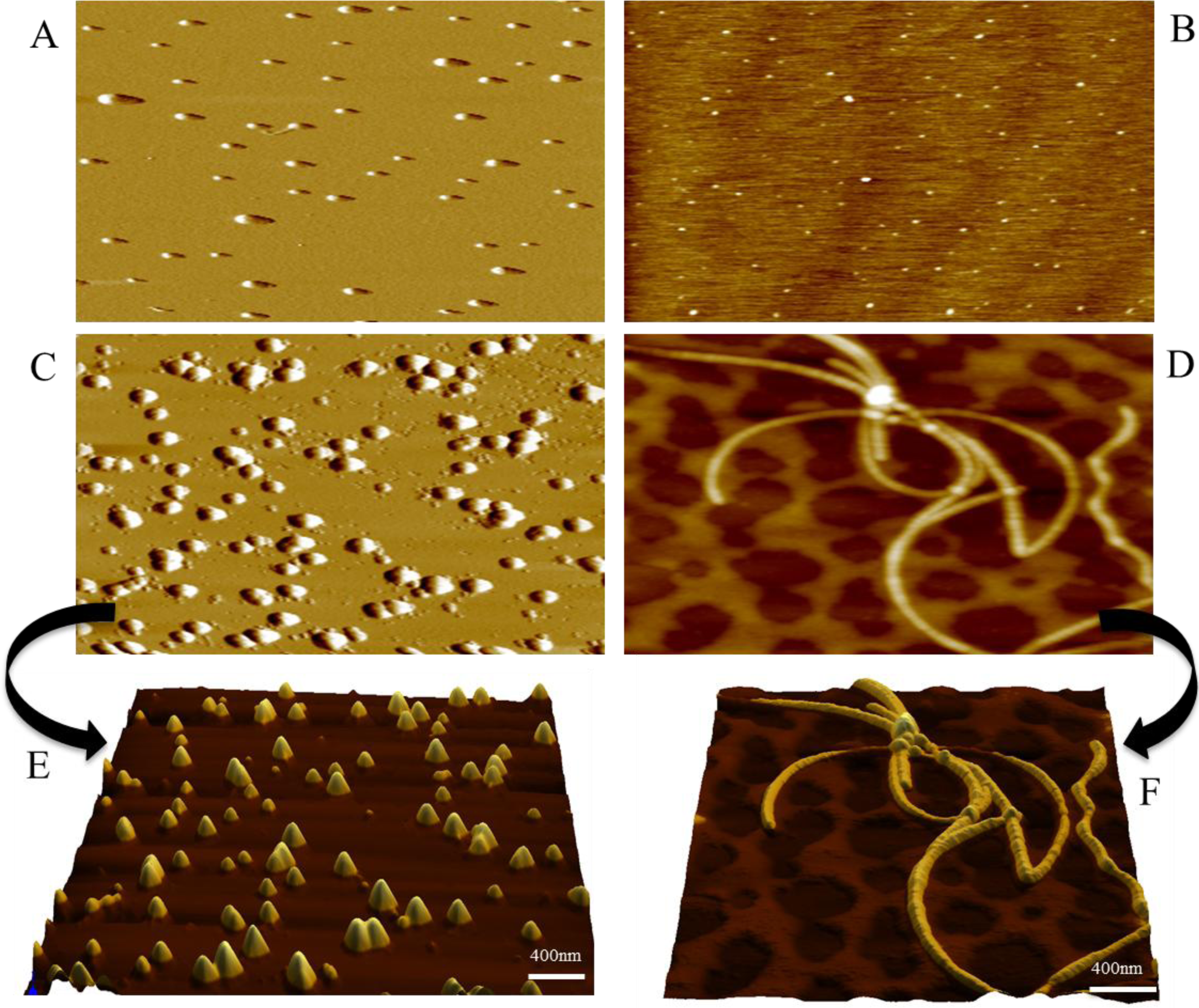
AFM images depicting morphological features of natural and induced oligomers of aS. Panel A (left) shows the natural oligomer of aS freshly prepared (0 h) and after incubation for 96 h (C) at 37 °C (750 rpm). Panel B shows oligomer form at 1 day of incubation of monomeric protein at 37 °C (750 rpm) and the bottom panel (D) depicts the fibril structure after 96 h of incubation. Initial height of the natural oligomers (NO) was ∼5nm and after incubation it increased to ∼22nm where as in incubated oligomers (IO) initial phase height was ∼1.5nm and after incubation it formed fiber of few micrometer in length. Figure C and D show 3D image of natural oligomer and fiber species after 96h of incubation. Aliquots have been collected at particular time interval from incubated samples for imaging purpose. The images were recorded at room temperature 25 °C using particular protocol mentioned in the methodology section.

### Secondary structure and circular dichroism (CD) measurement

Intrigued by the surface morphology, further assessed the secondary structural feature of the protein for its behavioural anomalies of the two different oligomers. Although the formation of aS fibrill is a condition dependent manner but these oligomers are characterized by their intertwined structures. aS is an intrinsically disorder protein and it prefers to remain as a disordered conformation when it is in monomeric form as indicated by negative elipticity marked at 198 nm (Figure 2). However, its structure altered depending upon it states of aggregation. Figure 2 represents CD spectrum of the two types of oligomers. The oligomeric species formed by lyophilisation (NO) showed negative maxima at ∼201nm and the conventional oligomeic species (IO) showed negative peak at ∼197 nm as reported in earlier studies^43^. However, the CD signature was quite broader in natural oligomers. Generally 195 and 198 signatures come due to electronic transition between π—π^*^ and the longer wavelength feature arises due to n—π^*^. In soluble forms of the protein, the negative signatures suggested the random forms as the β-II type^44^ conformation. The oligomeric may be enriched with β-II type formation most probably located within interior surface of oligomers. All these kind of conformation was further validated with Raman signature of the oligomeric structure. Induced oligomers also showed slightly negative trend at ∼ 218 nm. As the incubation continued at 37°C the induced oligomers (IO) gradually converted itself to fibrillar form and showed the negative peak at ∼218nm which suggested that the transformation of the protein’s secondary structure to β-sheet structure. Interestingly, NO did not change its conformation (Figure S2) throughout the incubation period. These results once again confirm that the natural oligomeric species are stable in nature than the induced one.

**Figure 2:**
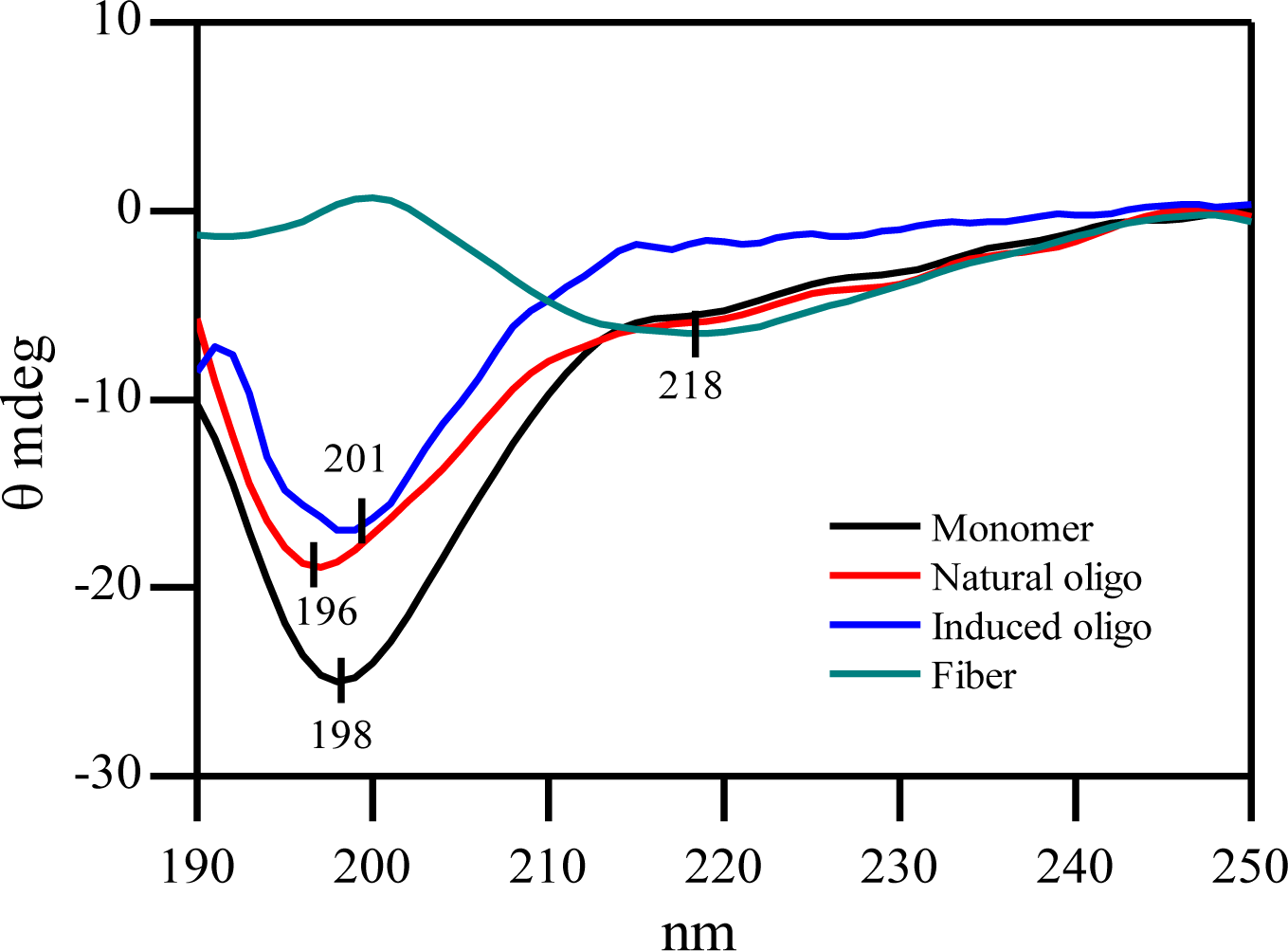
Circular Dichroism (CD) spectra of monomer, induced oligomer and natural oligomer in 20 mM sodium phosphate buffer, pH-7.4. The path length of the measuring cell was 1 mm. The sample preparation details were described in material method section.

### Probing the hydrophobic surface exposure

Existence of exposed hydrophobic surfaces is routinely determined by measuring the fluorescence of ANS bound to the hydrophobic protein surfaces. ANS preferably binds to the exposed hydrophobic surface of protein and produces significant fluorescence. In order to assess the exposer or increase of protein hydrophobic surface ANS fluorescence was measured separately in the presence of both type of oligomeric aS aggregates. In the presence of natural oligomers ANS showed no significant amount of fluorescence enhancement (Figure 4) and it was similar to the ANS in aqueous buffer solution. However, induced oligomer showed higher intensity of ANS fluorescence compared to the ANS in buffer solution. The lower panels in Figure 4 displays the ANS fluorescence intensity with aS oligomers at time point of incubation. ANS intensity in the presence of NO did not alter significantly over time of incubation whereas in IO it gradually increased over span of time. This suggested that natural oligomers contained lesser amount of hydrophobic surface compared to induced oligomer to bind ANS. In the induced oligomers, possibly the hydrophobic collapse extended to initiate the formation of cross beta sheet structure formation this hydrophobic surfaces are now capable to entrap the ANS molecules. The hydrophobic surface remains exposed on the outer surface.

**Figure 3:**
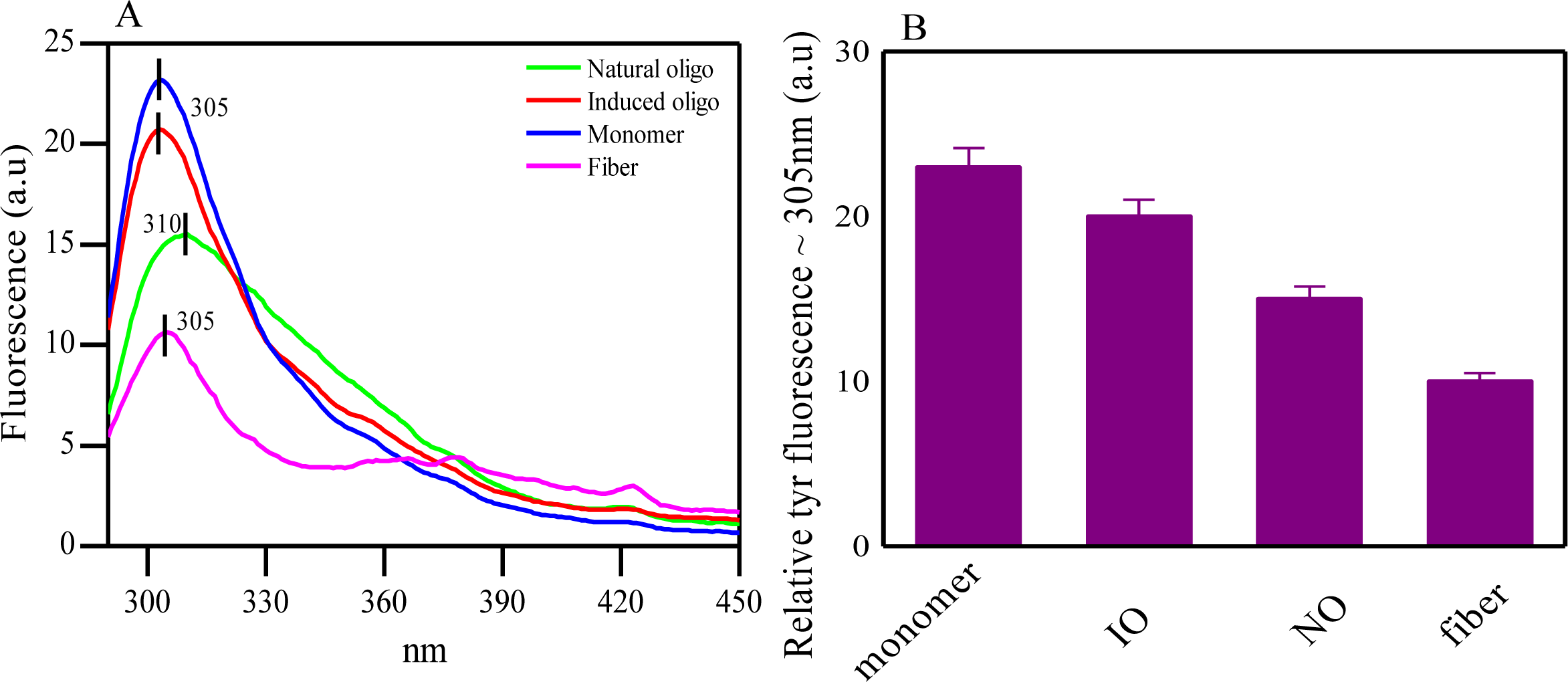
(A) Intrinsic tyrosine fluorescence of aS monomer, natural oligomer, induced oligomer and fibrillar aggregates. Fluorescence were measured upon excitation at 276 nm and slit width were kept at 5 nm. Fluorescence spectra were collected in 20 mM sodium phosphate buffer, pH-7.4. Right panel (B) depicts relative tyrosine fluorescence of the collected spectra at ∼ 305nm.

**Figure 4:**
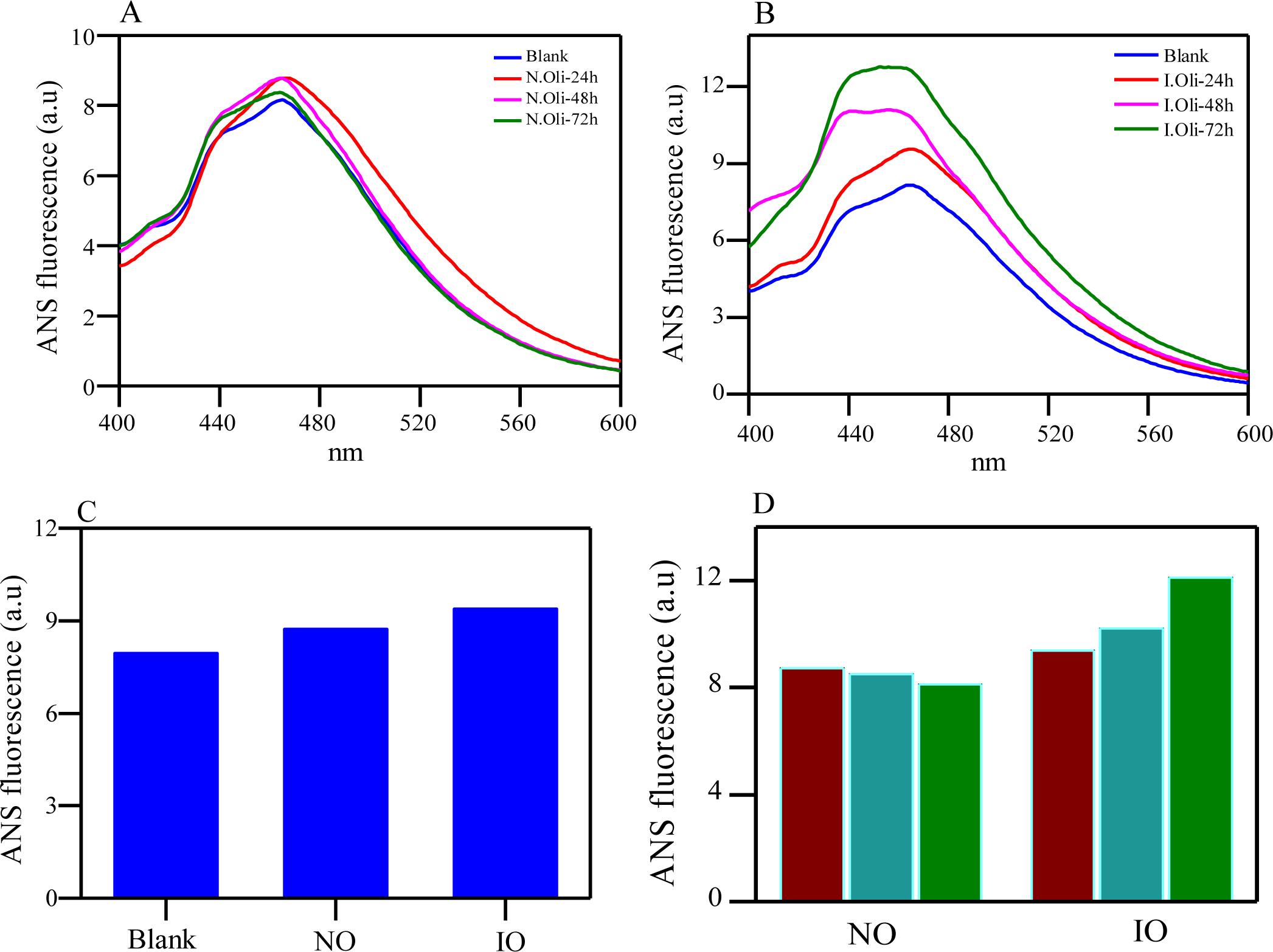
ANS fluorescence in the presence of aS monomer and other assembly structures. Upper panel showing fluorescence spectra of ANS alone and along with natural oligomer (A) and induced oligomer (B) were captured at particular time interval from the aged samples under particular condition (mentioned in the material method section). Lower panel (C) showing ANS intensity at initial stages of incubation. Right pane (D) showing changes in ANS intensity of oligomers over different time interval (magenta-24h, cyan-48h, green-72h). In natural oligomer ANS intensity remained unchanged throughout the incubation period. Samples were prepared according to described methods and spectra were recorded using 10 mm path length quartz cuvette at room temperature 26 °C.

The intrinsic fluorescence from the tyrosine residue has been used in several earlier studies to explore the interaction of peptides and proteins and their aggregation states.^45^ Different panels in Figure 3 show the tyrosine fluorescence of aS monomer, oligomer and fibrillar aggregates. Natural oligomer shows broader fluorescence band at ∼310 nm. However, fluorescence peak was much sharper for the monomer, induced oligomer and fibrils. It was also evidenced in the intrinsic tyrosine fluorescence behaviour observed higher Tyr fluorescence of induced oligomeric species compared to the natural oligomer. It signifies strong interactions among the units of natural oligomers. Monomeric form remains highly random configuration and as a result its exposed Tyr gives higher fluorescence than oligomeric conformations (Figure 3). All these were further confirmed by thioflavin (ThT) fluorescence assay.

### Structural elevation vs retardation

To investigate whether there is any kind of structural alteration (elevation to β-sheet structure) present/and occurring during the incubation period, thioflavin T (ThT) fluoresence assay was performed. ThT is a small molecular dye that becomes highly emissive upon binding to β-sheet rich amyloid protein fibrils. Figure 5B and 5D shows that natural oligomer caused no enhancement of ThT fluorescence and maintained similar level of fluorescence. Thus, throughout the incubation period natural oligomers retained its conformational state unaltered and ThT dye failed to bind effectively with these oligomers even after longer incubation of time. As a result there was no increment in fluorescence signal. ThT binds to the exposed β sheet and non-β sheet cavities with a diameter of 8-9 A^° 46^ of the proteins and if these β is located in the interior of the proteins it could not interact. Thus fibril formation kinetics was made by measuring ThT fluorescence at different time point of incubation. A gradual increment in the fluorescence intensity was observed for incubated induced oligomers. After 48h of incubation ThT of IO increased significantly. Figure 5A and 5C illustrated changes in ThT intensity upon interacting with induced oligomers over the time period. These oligomers showed typical invitro aggregation kinetics with sigmoidal curve in its nature (Figure 5E). The lag phase persists upto 24h followed by a growth phase of 80h. During the growth phase oligomers rapidly undergo through conformational changes and attain fibrillar morphology. Structural elevation of these oligomers indicates its transient nature in this time period. Natural oligomers (NO) showed very little increment of ThT fluorescene as shown in the Figure 5F. However after 50h of incubation of the natural oligomer small increment in ThT fluorescence was noticed which might be attributed due to the formation of non-fibrillar small aggregates possessing some affinity to the dye. Thus NO showed relatively higher stability. This indicates that the oligomers formed via incubation gradually altered its conformation to β-sheet rich amyloid fibril; However, the natural oligomers retained its oligomeric conformation throughout the incubation period.

**Figure 5:**
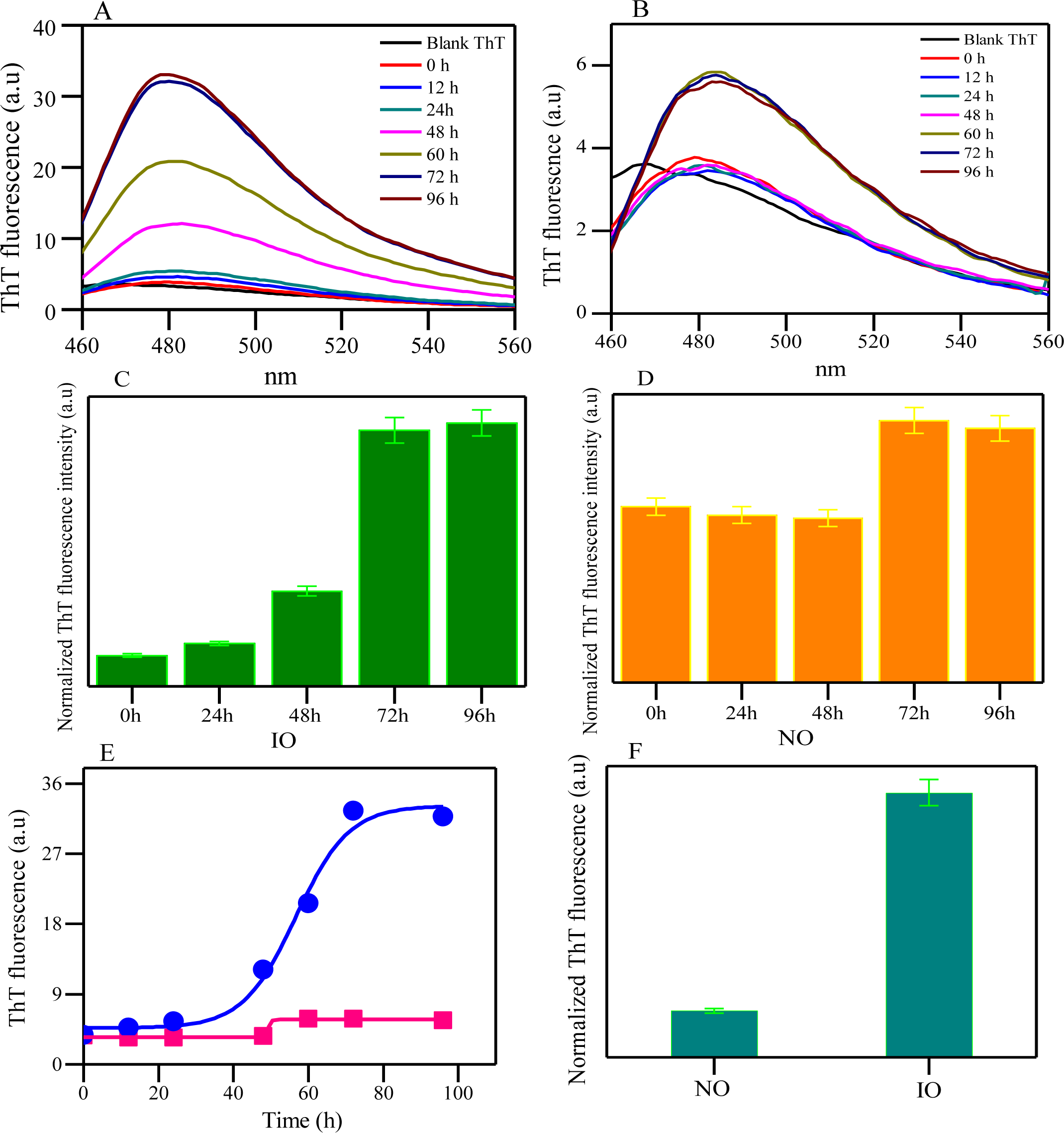
ThT measurement for determining growth kinetics of oligomeric species. Figure A and C denote changes in ThT intensity in presence of Induced oligomer over incubation time period. Figure B and D indicates changes in ThT fluorescence over incubation period of natural oligomers. In Figure E Blue line indicates typical amyloid fibrillation pattern of induced oligomer whereas Pink line indicates no fibrillation of natural oligomeric species. Samples were prepared according to described methods and spectra were recorded using 10 mm path length quartz cuvette at room temperature 26 °C. Natural oligomer preferred to retain its oligomeric form throughout the incubation period. Lower panel F shows changes in fluorescence intensity of ThT of the both oligomer at equilibrium stage.

### Raman signature and structural variation of two type oligomers

Raman Spectroscopy provides crucial insight of secondary structure of protein assembly and different form of its aggregates. Figure 6A and 6B display the Raman spectra in the region of 600-1800 cm^-1^ of natural and induced oligomers, respectively. The induced oligomers obtained by incubation monomeric sample for one day and the natural oligomers are prepared from the stored monomeric sample as discussed in the materials and method sections. The vibrational modes and Raman peak assignments are listed and given in Table S1. Natural oligomer (NO) produced a amide I vibration band at 1669 cm^-1^ and one small shoulder band at 1615 cm^-1^ attributed to the ring mode vibration of tyrosine residues. The peak width at half-height (PWHH) of the band was ∼46 cm^-1^. The band position was similar to monomeric protein solution (Figure S3).^47^ The symmetric nature and the significant band width indicated the presence of multiple component of protein secondary structure. The hump at ∼1654 cm^-1^ (Figures 6F) indicated the presence of globular fold with α-helical structure.^39,48–52^. Induced oligomers (IO) showed a similar Raman band patterns, however the amide I region was sharper and centred at 1668 cm^-1^ with PWHH ∼36 cm^-1^. Amide I region of NO is comparatively broader than the induced oligomeric species which indicates its more heterogeneous nature.

**Figure 6:**
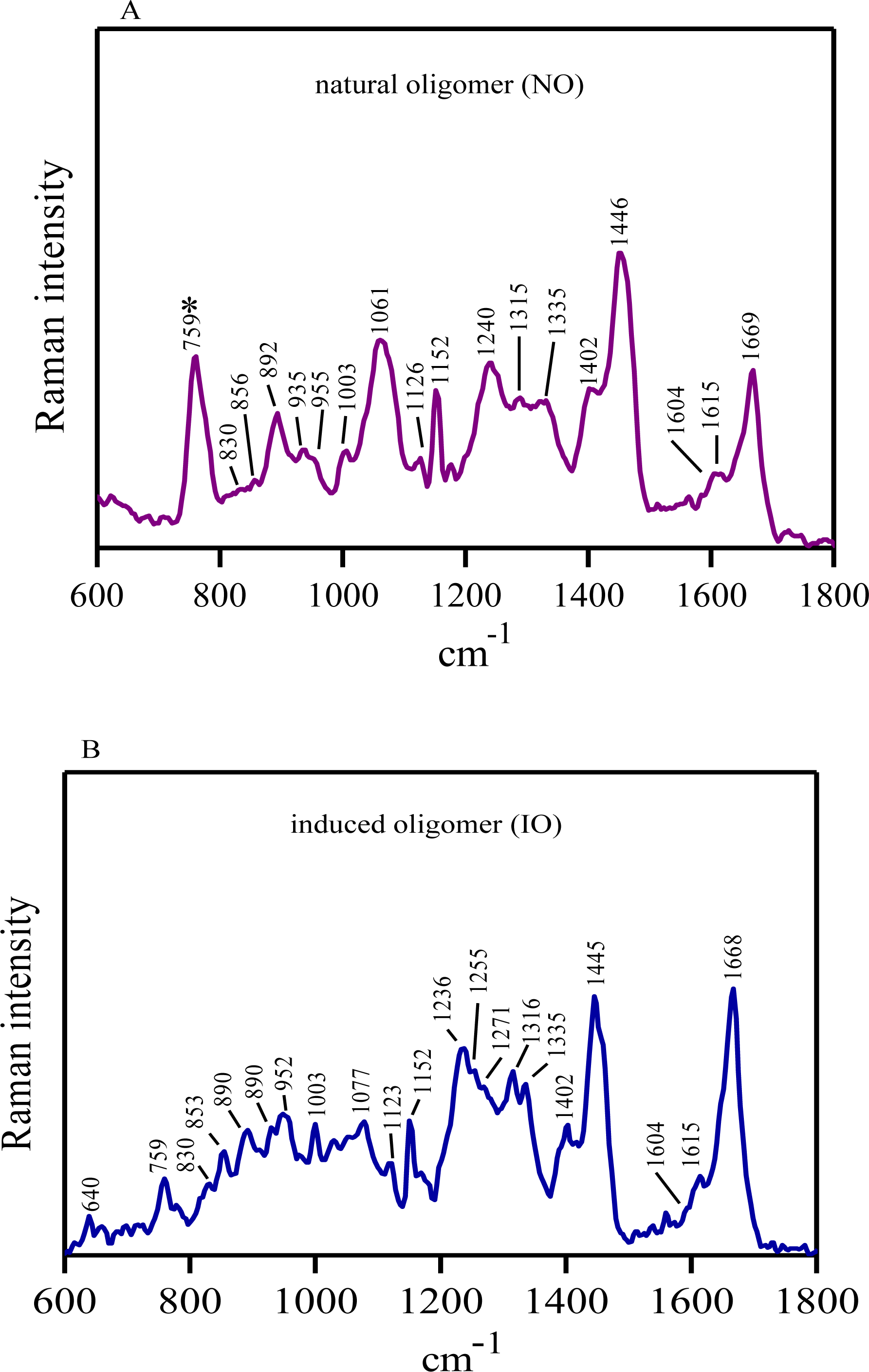
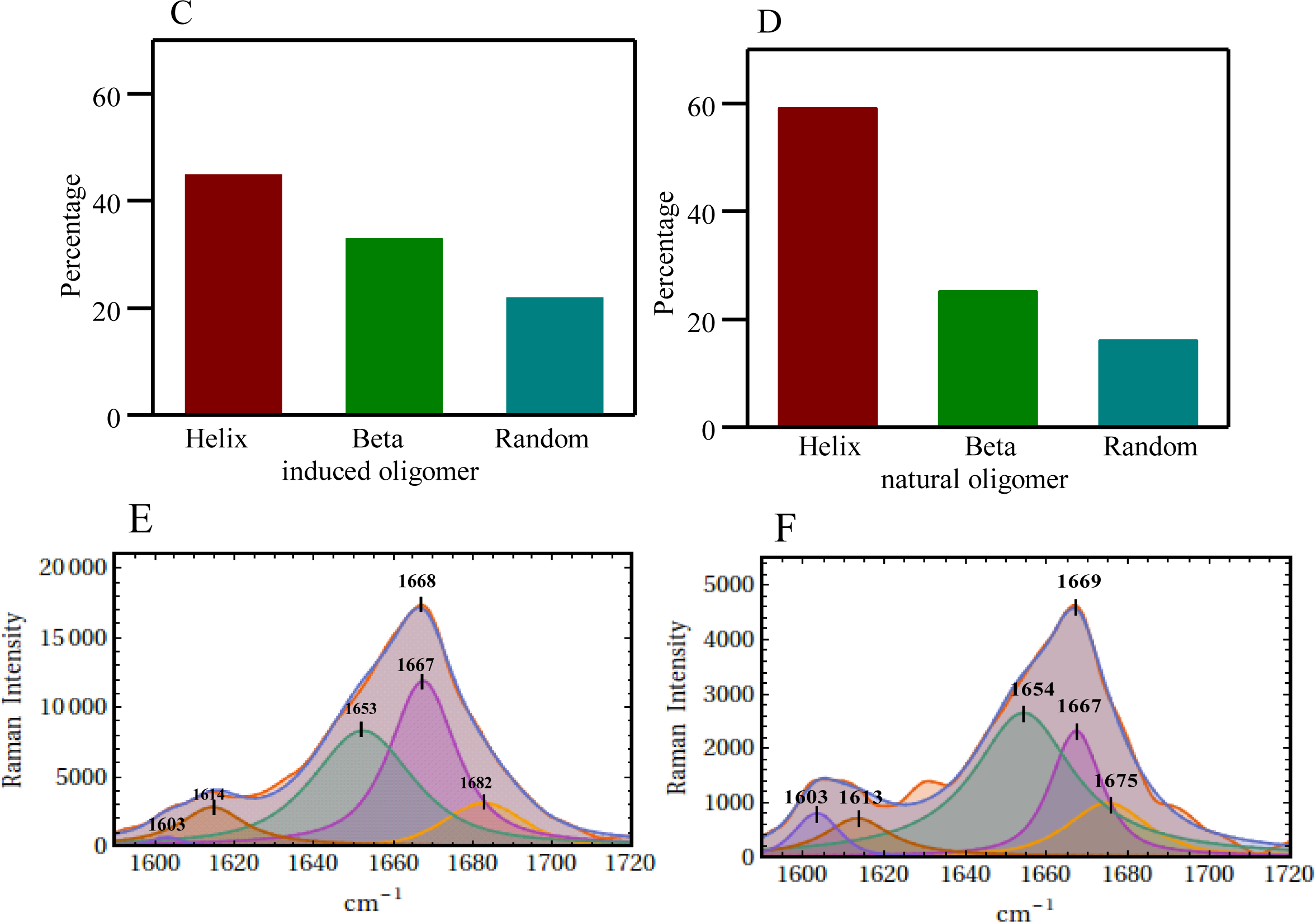
Raman spectra of oligomer in the frequency of 600-1800 cm^-1^ by 632 nm Laser. Oligomer solution (200µM) was prepared in 20 mM sodium phosphate buffer pH-7.4 and it was drop casted on a glass cover slip and spectra was recorded at room temperature (25°C) after air dried. Laser power at the source was 20 mW and ∼2 mW at sample. Figure (A and B) shows Raman spectra of natural oligomer and induced oligomer respectively. Figure (E and F) denotes curve fitting analysis of Amide I region (1590–1720 cm^-1^) of induced and natural aS oligomers. The yellow and deep blue line indicates the experimental and fitted spectral signature respectively. Three component bands of the amide I regions are shown in magenta, green, yellow shades. The fitted peak positions of the assigned aromatic residues are also indicated in violet and brown shades. Figure (C and D) indicates secondary structure of both the oligomers derived from band fitting analysis. *represents buffer contribution.

The amide-III region which involves in-phase combination of N-H deformation and C-N stretching shows overlapping bands in 1230-1300 cm^-1^ range. Both the oligomers exhibit bands at ∼ 1252 and ∼ 1242 cm^-1^. The amide-III band at 1255 cm^-1^ is assigned to mainly disordered or random conformation within the protein. The ∼1240 cm^-1^ band corresponds to distorted β-sheet/ β-strand structure. In the induced oligomer the band position at ∼1242 cm^-1^ shifted slightly to lower frequency region of 1236 cm^-1^ and indicated slight increase β-sheet conformation. These signatures indicated simultaneous coexistence of disordered state as a part of folded conformation in the overall secondary structure of the protein in oligomers. Earlier reports characterized that Raman signature near ∼1340 cm^-1^ indicates vibration involving C_α_-H bending and C_α_-C stretching. Although, the band near 1345cm^-1^ is outside the range attributed to the amide III region in proteins. However, it could be emerged due to the side chain contribution of Lys, Gly and Ser. In case of natural oligomeric species a signature band comes at 1335 cm^-1^ which is the signature of α-helical conformation and stretching of C-N bond of proteins^53^. It was interesting to note that the Raman intensity at ∼1335 cm^-1^ started weakening, though separated well with the 1316 cm^-1^ band, in induced oligomeric form. It was substantially decreased in the fibrillar state (Figure S3). It signifies enormous reorientation as well as decrease in the helical structure of the proteins and it may exist as a helical kink connected via short loops of few amino acid residues. This structure is highly up holded with the capsid protein of virus^54^.

The region between 800-1200 cm^-1^ gets many signature band 955 and 935 cm^-1^ which denotes the presence of backbone tertiary signature of helics^55,56^. This also corroborates a model like viral procapsid^57^in which folding of α helical structure is essential for assembly of natural oligomer and appeared at 930-940 cm^-1^. In NO the stretching peaks centred at ∼935 cm^-1^ remained stronger and indicated large content of helical compared to induced oligomers and fibrils. The deformation of CH_2_ is well indicated by 1316 cm^-1^ but it is relatively diminished in case of natural oligomeric species. Furthermore the deformation of CH_2_/CH_3_ group could be noted here by abroad spectra of 1445cm^-1^.One other possibility that could also attribute to this phenomena is the carbonyl stretching of the amino acid proline^58^. The CH_2_ deformation band got shifted to 1315 cm^-1^ with less pronounced shoulder than the induced oligomers.

### Curve fitting to amide I band

In the Raman spectra amide-I mode of vibration appears in 1630-1700 cm^-1^ region and it arises arises mainly due to the carbonyl stretch of the polyamide backbone of the protein. It is also contributed by vibration of out of phase C-N motion. The band is often used to accesses the secondary structure in different physical states of protein. The amide I band of the oligomers were broad, indicating a mixture of conformations and a simple curve-fitting technique was applied to measure the contributions from different secondary conformations. The secondary structural components were mentioned in the Table 1. The amide-I region centre at 1669 cm^-1^ and it’s PWHM was measured to be ∼ 46 cm^-1^. The component band at 1654 cm^-1^ indicates its helical conformation and ∼59% residues (Figure 6F & 6D) preferred this structure; it was comparatively higher than the induced oligomers. However, the β-sheet/strand components were measured to be 25%. It is lower in amount than the induced oligomers. Interestingly, the PP-II component (1675cm^-1^) was also less than the induced one. It was restricted to ∼16% only. Major signature of the Amide-I band towards 1669 cm^-1^ denotes extended β conformation. It produced a quiet similar structure of antiparallel pleated β sheets found in silk fiber of *Nephila clavipes*^59^.

**Table 1.** Band fitting of the amide I band profile of natural and induced oligomer of aS.

| Peak no. | Center ($\text{cm}^{-1}$) | Width (nm) | % Area |
| --- | --- | --- | --- |
| **Natural oligomer** |  |  |  |
| 1 | 1654 | 29 | 59 |
| 2 | 1667 | 15 | 25 |
| 3 | 1675 | 23 | 16 |
| **Induced oligomer** |  |  |  |
| 1 | 1653 | 29 | 45 |
| 2 | 1667 | 20 | 33 |
| 3 | 1682 | 23 | 22 |

In induced oligomeric species (Figure 6C & 6E) the helical presence was 45% (band component 1654 cm^-1^), β-sheet/strand was ∼ 33%. β-component was represented by the fitted component band 1667 cm^-1^. PP-II like conformation was 22% as indicated by the component band 1682 cm^-1)^. Two additional components were due to ring mode vibration of the aromatic ring of Phe (1603 cm^-1^) and Tyr (1614 cm^-1^). The appearance of a strong band at 1668 cm^-1^ indicates strong hydrogen bonds with in the β structure and could be antiparallel. Here the conformational distortions most likely aroused due to the disparity in the arrangement and its interaction with side chains of amino acids during packing. However, its structure could be assigned to PG-II conformation^59^.

### Cellular Toxicity

There has been considerable curiosity towards the toxicity of aS oligomers. Several studies well demonstrate that the oligomers were highly toxic compared to fibrills^60^. Therefore, intrigued to determine the toxicity of the two different oligomeric species. Neuronal SH-SY5Y cells were treated with the oligomeric species of different concentration in a time dependent manner. In comparison cells treated with natural oligomeric species showed higher toxicity compared to the induced oligomeric species. However, cells treated with monomeric aS did not exhibit any toxicity. Primarily these results indicate that availability of cellular level of aS oligomers can influence the toxicity of the cellular system. From Figure 7 it is evident that the natural oligomeric species at concentration of 10μM exhibited higher toxicity compared to the induced oligomeric species and it is also capable of inhibiting the proliferation (Figure 7) of neuronal cells. Earlier reports indicated that the spherical oligomers of Aβ pertide possessed the toxic moiety rather than its aggregates or fibrillar forms^61^. In a concurrence found that the nonfibrillar oligomers of aS are capable of disrupting the membrane integrity of neuronal cells more than the induced oligomeric species. The association of aS with vesicles is regulated by synaptic activity where it detaches from the vesicles after stimulation of neurons and then it reassociates.^62^ Because of binding of aS to synaptic vesicles it is possible that transmission from synapse could be directly or indirectly a potential target of synuclein toxicity^63^. Natural oligomeric species bears higher amount of antiparallel β sheet compared to the Induced oligomers as found from CD analysis, this may be the crucial player for exerting toxicity. These nonfibrillar soluble oligomeric species comprised of disordered monomeric aS species holds significant amount of β-Sheet structure in its interior which is evident from Raman signature. The structural property and morphology of the two different species of oligomers affected the cellular toxic nature in a dramatic fashion.

**Figure 7:**
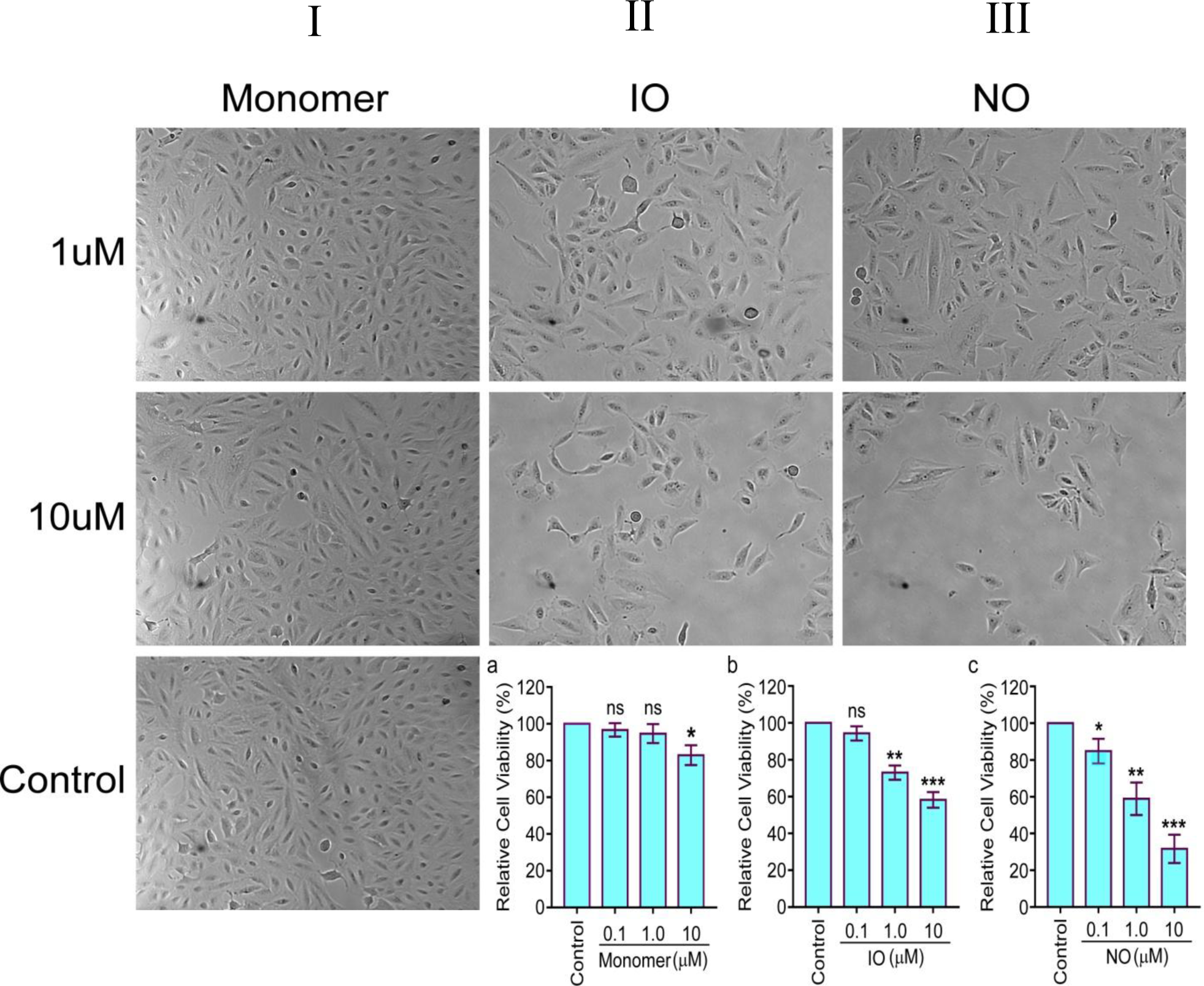
Cytotoxicity assay - microscopic images of SH-SY5Y cells after 72 hours treatment of different doses of (I) monomer, (II) induced oligomer (IO), (III) natural oligomer (NO) and monomer of aS. 1 µM and 10 µM (as indicated) protein concentration were used for treatment of neuronal cell lines. Images were captured at 10X objective. From the figure it is evident that NO can prohibit cellular proliferation with higher efficacy than the IO. Effect of Monomer, IO and NO on proliferation of SH-SY5Y neuroblastoma cells were measured by MTT assay (Figure a, b & c respectively). Cells were treated with different concentration peptide (NO, IO and Monomer) and incubated for 72 h in a 5% CO2 incubator followed by MTT assay was performed. Natural oligomeric species damaged the cell to a higher extent.

### Oligomer stability, toxicity and model

Protein folding is a process of spontaneous molecular self-organisation through which after different molecular ensembles protein passes its way to folded structure.^64^ The free energies associated with this ensemble structure define its molecular configuration. The essence of process of organisation is that it begins with many possible states and it terminates with few states. The energy landscape theory postulates that there is existence of several scenarios during the folding and aggregation of proteins (Figure 8). In some of the folding scenarios the kinetically crucial stages of folding occurs in presence of large density of configurations. The monomeric form of intrinsically disordered proteins is kinetically unstable and it can under goes through two different path where one form simply interact among themselves in such a way that it becomes kinetically trapped to particular stable form. This form does not interact further and they are the natural oligomers. Whereas in other scenario monomer interacts to form transient unstable oligomer which further interacts to form relatively more stable fiber. The energy level of this fiber configuration is very low and they are stable in nature. The induced oligomeric forms are less toxic compared to natural oligomeric form. This denotes that energy level in course of protein aggregation could be a key modulator behind altering level of toxicity. Thus natural oligomer, as supported by several observations, trapped with helical content to energetically stable state compared to induced oligomers which pass through a small energy barrier and transformed into β-sheet rich fibrillar aggregate.The seeding of β-sheet conformation and hydrophobic zipping may initiate in its early oligomeric state itself. However, monomeric units in natural oligomer, may be intertwined with different hydrogen bonding network that provide both the stability and caused retardation in fibril formation.

**Figure 8:**
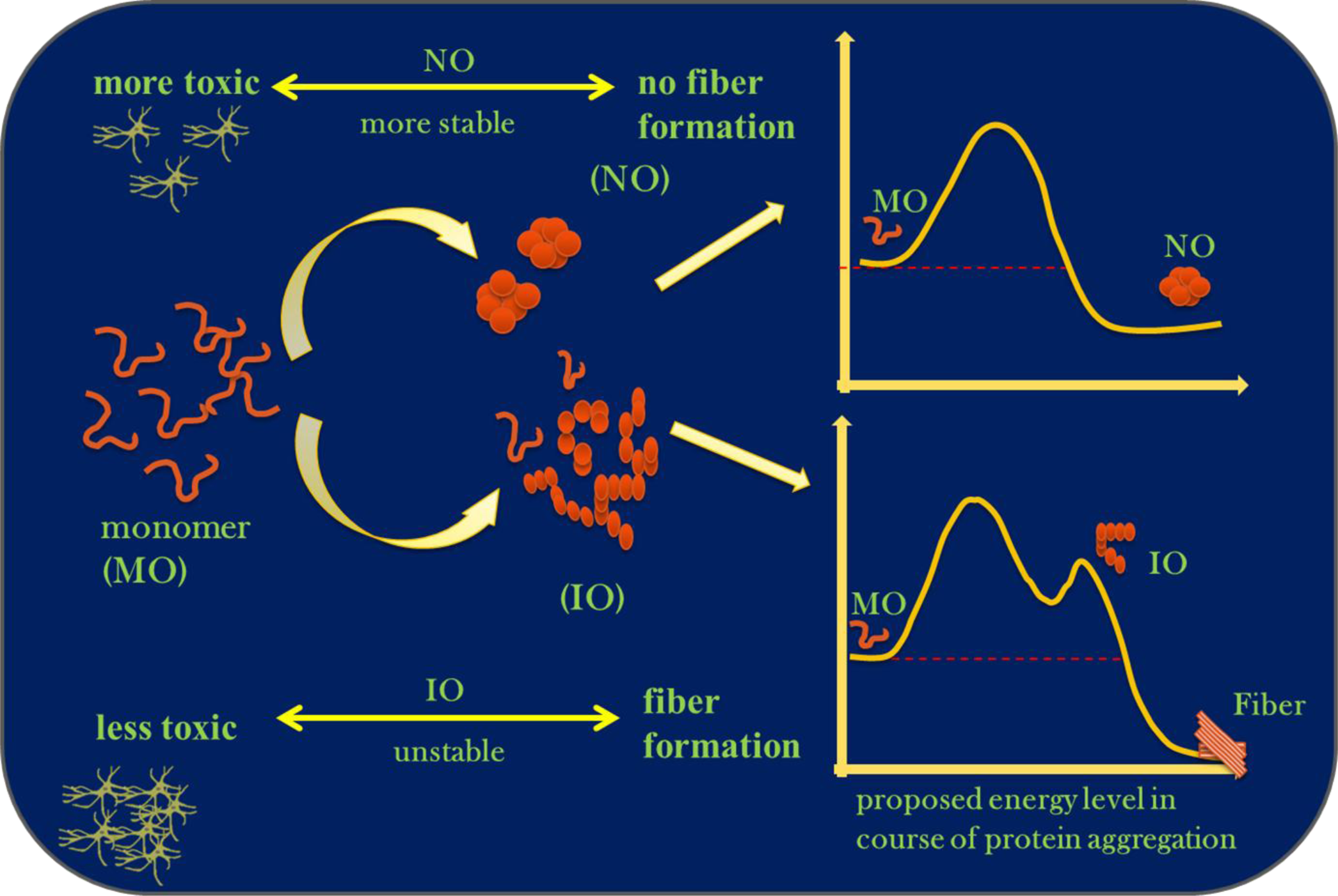
Model showing proposed energy level followed in course of aggregation by oligomers. Monomer aS interacts among themselves in such a way that it forms two types of oligomer NO and IO. NO is quiet stable and it did not form fiber whereas IO being unstable interacts further and form fiber species.

## Conclusion

The novel aS oligomer characterized here by its structure and toxic level could provide fundamental understanding for developing new therapeutics against Parkinson disease (PD). All the information accrued by the comparative studies indicates natural oligomer of aS possess morphological similarity with the induced oligomers but it has some structural difference that makes it relatively stable and toxic than the induced oligomers. Emphatically the surface hydrophobicity and exposed β content was comparatively lower in the natural oligomeric form. It could be the main reason behind its non fibrillar form and structural retardation to evolve further to β-sheet rich amyloid like fibrillar aggregates. The induced oligomer showed its higher seeding capacity and forms amyloids fibrills very rapidly with a sigmoidal growth curve.

From the point of structural differentiation of natural oligomer from the induced oligomeic the former one showed more heterogeneous structure with distorted β sheet located within its interior region. However, it showed an intermingled structure of helices as well as β sheet. Thus the secondary structure of the oligomer is different from the conventional fibrill forming oligomers. Similar to the membrane fusion by viral protein the helical content with in the natural oligomeric species may augment the disruption of cell during its transmission^55^. Several factors at the interior and exterior can initiate partial or complete folding of the disorder protein aS such as lipid, membrane, ions and macromolecular crowding^65,66^.In our study the nonfibrillar oligomers are found to be stable in their conformation and these soluble oligomers comprised of ensemble structure between helices and distorted β sheet. These can play crucial role in the steps of aggregation and these should be taken into account while targeting for therapeutics. Future study regarding the mechanism by which natural oligomeric species exacerbating the toxicity within the neuronal cell line will be interesting. Natural oligomers with helical preferences may interact well with cell membrane and become more toxic to neuronal cells.

## Material and Methods

### Expression and Purification of α-Syn

The plasmid containing aS gene was a kind of gift from HilalLashuel (Addgene plasmid # 36046)^67^. Then aS was expressed in *E*.*Coli* BL21(DE3) using plasmid pT7-7 encoded with wild type human aS gene. Followed by expression and purification was done by earlier established protocol. First bacterial cells were grown using LuriaBroth media in the presence of antibiotic amphicillin(100µg/ml) as a selection agent. Then expression of aS was done by the induction of 1mM Isopropyl-β-D-thiogalactopyranoside (IPTG). After that bacterial pellets were harvested by centrifugation at 5000 rpm for 15 min at 4°C.Then the pellets were resuspended in lysis buffer containing 30mM Tris pH 7.5, 1mM ethylene diaminetetra acetate, 1mM phenylmethanesulfonyl fluoride, 1mM dithiothreitol. Followed by cells were vortexed to dissolve the pellet completely until no clumps were visible. The lysate was centrifuged at 18000 rpm and supernatant was collected. Then it was subjected to acid precipitation until it reached pH-4.This step helps in removal of undesired proteins. Again it was centrifuged at 18000 rpm and supernatant was collected for overnight dialysis in 50mM Tris and 300mM NaCl. After that ion exchange and size exclusion chromatography was performed to separate the protein. The relevant fraction of the protein was collected and its purity was checked using SDS-PAGE and MALDI analysis (S1).

### Sample Preparation for Spectroscopy

Spectroscopic studies were performed by dissolving lyophilized (stored at −20°C for 7days) aS in 50mM Tris-HCl pH-7.5 and it was centrifuged at 15000 rpm for 5 min to remove any insoluble particles and loaded into a Hi load 16/600 Superdex 200pg, GF column which was equilibrated with buffer. Then relevant fraction of the protein (Natural Oligomer & monomer) was collected (Figure S1) and concentrated using a Millipore Amicon Ultra 3Kd cut off concentrator. Protein concentration was checked by UV absorbance measurement using extinction co-efficient of 5960M^-1^cm^-1^ at 276 nm. Purity of the aS protein was judged by performing SDS-PAGE containing a single band near 14Kd region. For both CD and Raman Spectroscopy studies 200µM of aS in 20mM sodium phosphate buffer was used. Samples incubation was done in following way - 200µM of aS was dissolved in 20mM Sodium Phosphate buffer and incubated under constant shaking 750rpm at 37°C.

### Circular Dichroism study

Far UV-CD measurements were recorded on a model no. J-815 spectrophotometer (JASCO) using a 1mm pathlength cuvette at 25°C temperature. Spectra were acquired with a scan speed of 50nm/min with an interval of 2sec. For each sample three scans were done and the buffer background was subtracted. 200µM of aS was dissolved in 20mM Sodium Phosphate buffer and incubated under constant shaking 750rpm at 37°C. For CD analysis 10µl of the sample was dissolved in 200µl of the 20mM sodium phosphate buffer from incubated sample at a particular time interval to get a diluted solution of a concentration about 10µm.

### ANS-Binding Fluorescence Assay

A stock of ANS (1-Anilino-8 Napthalene Sulfonate) dye of 1mM was prepared in double distilled water. To determine the hydrophobic exposure in oligomer 500µl of assay buffer containing 25µM ANS in 20mM Sodium Phosphate pH-7.4 was mixed with 3µM of (500µM) natural oligomeric solution. It was mixed properly before fluorescence experiments. The mixed solution was then taken in a 1cm path length quartz cuvette and the fluorescence measurement was performed using a PTI fluorescence spectrophotometer. Emission spectra were acquired from 400 nm to 600 nm with excitation at 370nm and the integration time was 1 sec. Slit widths were kept at 5nm each during measurement.

### ThioflavinTAssay

For ThT fluorescence measurement a stock of 5mM ThT was made and fluorescence measurements were performed using a 10mm path length quartz cuvette. To measure amyloid fibrillation Oligomeric solution of aS mixed thoroughly with 500µl of 20mM Sodium phosphate buffer pH-7.4 to obtain a solution of 3µM α-Syn. Then solution of ThT from 5mM stock was added to it for obtaining 20µM and mixed thoroughly for acquisition of fluorescence emission spectra. Fluorescence measurements were obtained using PTI fluorescence spectrophotometer from 450-560nm with excitation at 440nm and slits were kept at 5nm.ThT fluorescence peak intensity at 482 nm was plotted against time, analysed and fitted to the sigmoidal curve using equation^68^.

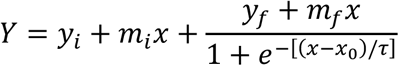

Where Y is the ThT fluorescence intensity at particular time *(x), x* is the incubation time, and *x*_*0*_ is the time to reach 50 % of maximal fluorescence; other parameters are determined by the fitting. The lag time is defined by *x*_*0*_−*2τ*. Apparent rate constant (1/ τ), *m*_*i*_ and *m*_*f*_ are two constants (linear coefficients).

### AFM Study

aS solution were taken out at particular time interval for AFM analysis from the incubated sample. The aliquot was casted to a freshly cleaned muscovite mica substrate and kept at room temperature for 30 sec. The mica surface was then rinsed with Millipore filtered water twice (100µl) to remove loosely bound protein and then it was dried. After that the sample was imaged immediately with tapping mode of AFM using a Pico plus 5500 AFM (Agilent Technologies, USA) with a piezoscanner having a maximum range of 9µM.The images were captured with a scan speed of 0.5 line/sec. Imaged were processed by flattening using picoview software.

### Raman Spectroscopy

Raman spectra were recorded using a STR Raman spectrometer which was equipped with a Olympus BX51 microscope and 500 mm focal length triple grating monochromator connected with CCD. Laser was focused through a 50× objective and the scattered light from the sample was collected through the same objectives. Oligomeric solutions of aS 10µl were first drop casted on glass slide and it was air dried for obtaining spectra. Samples were excited with a 630 nm wavelength laser light, with 30–35 mW of radiant power at the source and ∼2 mW at the sample. The recording time was 15 s, and number of scans was 50. The wavenumbers of the Raman spectra were calibrated with the Raman band of silica wafer at 519 cm^−1^. Spectra were recorded from a spectral range of 500–1800 cm^−1^. Spectra were processed with the GRAMS/A1 software.

### Cell proliferation assay (MTT assay)

For cell proliferation assay SHSY-5Y cells (2×10^4^ per well) were seeded in 12 well plates. After 72 h of treatment (Natural Oligomer-**NO**, Induced Oligomer-**IO** and **M**onomer at different doses), 1000 μl of MTT (0.5mg/ml) dissolved in phenol red free DMEM supplemented with sodium bicarbonate was added per well and incubated in a CO_2_ incubator for 3 h. The media was then discarded and 1 ml DMSO was added to it. After 10 min of incubation with DMSO the reading of the violet colour formazone was taken at 570 nm.

## ▪AUTHOR INFORMATION

**Corresponding Author**

Nakul C. Maiti

Phone: +91-33-2499-5940

Fax: +91-33-2473-5197

## Acknowledgement

Animesh Mondal thanks CSIR for the fellowship. Sandip Dolui thanks CSIR network project BSC0113, BSC0115 and BSC0121 for funding support. The authors thank T. Murganandan for recording AFM and also acknowledge other central instrumental facilities. The Raman instrument was purchased under DBT, New Delhi grant (GAP-299) to Dr. Nakul C. Maiti.

---

**Scheme 1:**
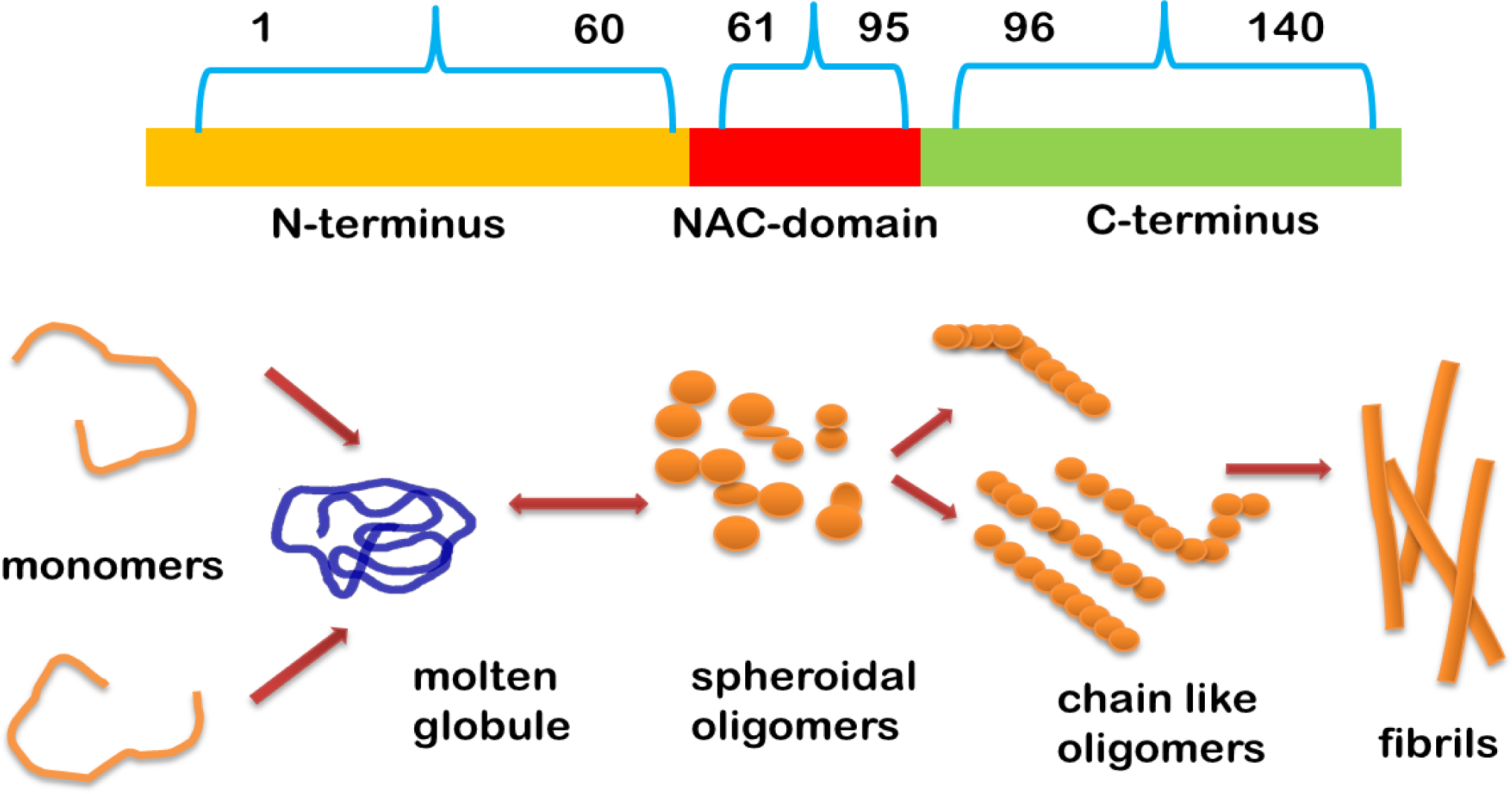
Model showing aS sequence region and conventional pathway of forming oligomers from monomer of aS. In conventional pathway monomer first under goes through molten globule conformation then it interacts further to form different morph of on pathway oligomers which will eventually form higher order fibrillar structure.

