## supplementary AM for "Structure Specific Neuro-toxicity of α-Synuclein Oligomer"

### Figures

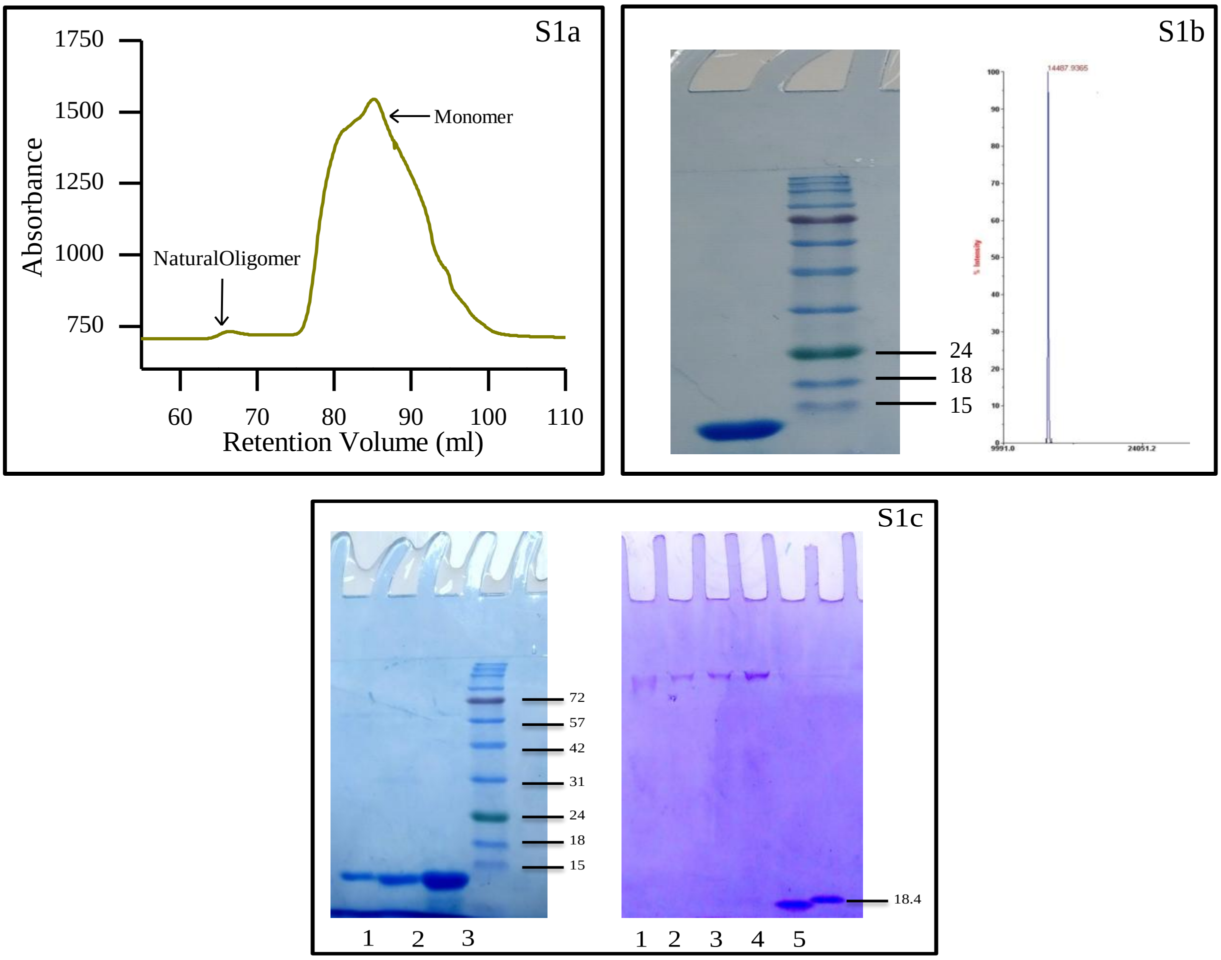


**Figure S1:** Upper panel (S1b) showing general sequence of αS and confirmation of aS by SDS-PAGE and sharp MALDI peak of purified αS. Upper left panel (S1a) depicts Seperation of Natural Oligomer and Monomer from lyophilized sample of aS after solubilization in Tris Buffer. Oligomeric samples were separated using Superdex 16/600 column (200pg). Oligomeric fraction was eluted at (65-70) ml region. Lower panel S1c (left) showed 15% SDS-PAGE of natural oligomer, induced oligomer and monomer of aS in lane 1,2 and 3 respectively. Lower right panel showed native PAGE of induced oligomer (lane1), natural oligomer (lane 2,3,4) and monomer (lane 5). For Native PAGE Lactoglobulin (18.4) was used as marker band. In Native PAGE both of the oligomers did not migrate and they remained in the stacking portion of gel due to their higher molecular weight. However in case of induced oligomer in the Native PAGE it appeared as smear.

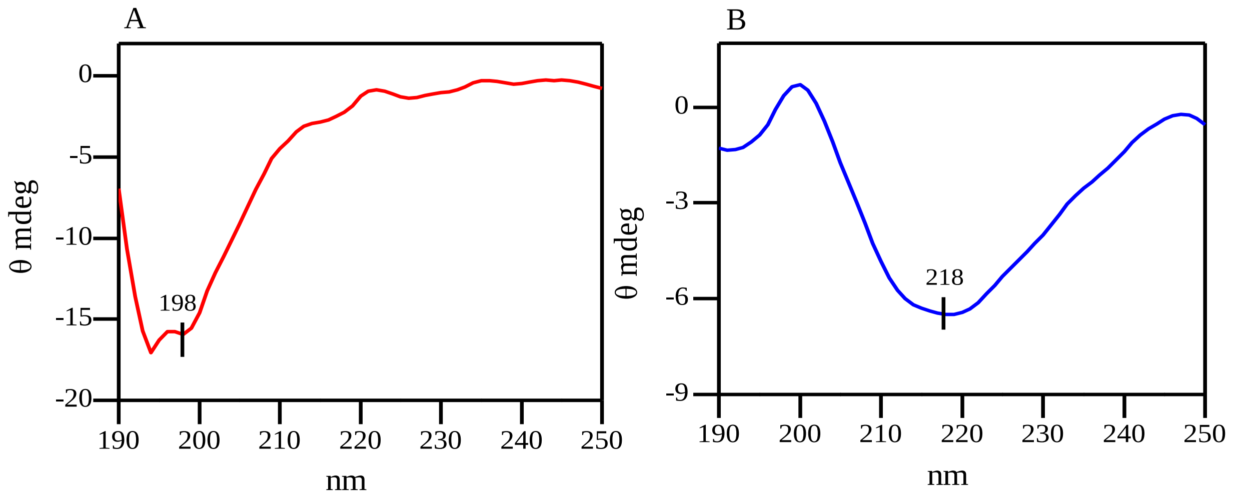


**Figure S2:** Left panel (A) showing CD spectra of natural oligomer (NO) and right panel showing spectra of induced oligomer (IO) after 4day of incubation (37˚C,750 rpm in 20mM sodium phosphate buffer.

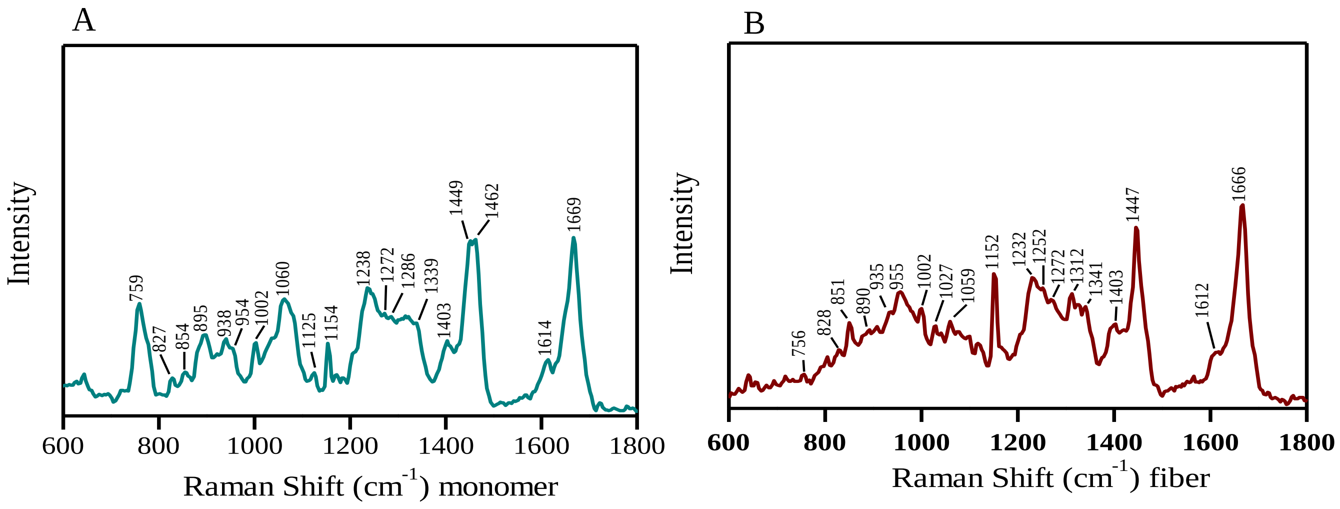


**Figure S3:** Raman spectra of aS monomer (A) and fiber (B) in the frequency of 600-1800 $Cm^{-1}$ by 632 nm Laser. Protein solution (200µM) was prepared in 20mM sodium phosphate buffer pH-7.4 and it was drop casted on a glass cover slip and spectra was recorded at room temperature ($25^{0}$C) after air dried. Laser power at the source was 20mW and ~2mW at sample.

**Table S1 :** Raman Vibrational bands ($cm^{-1}$) of natural and induced oligomer of aS.

| **Natural oligomer** | **Induced oligomer** | **Modes of Raman Vibration** |
|---|---|---|
| | 640 | Tyr |
| 830 | 830 | Tyr |
| 856 | 853 | Tyr |
| 892 | 890 | νCC |
| 935 | | α- helix |
| 955 | 952 | νCC |
| 1003 | 1003 | Phe |
| 1126 | 1123 | νCC |
| 1152 | 1152 | νCN |
| 1240 | 1236 | Amide-III,β-sheet |
| | 1255 | Amide III, random |
| | 1271 | Amide-III,α-sheet |
| 1315 | 1316 | CH2 deformation |
| 1335 | 1335 | CH2 deformation |
| 1402 | 1402 | symmetric νCO2 - |
| 1446 | 1445 | CH2, CH3 deformation |
| 1604 | 1604 | Phe |
| 1615 | 1615 | Tyr |
| | 1668 | Amide I, β-sheet |
| 1669 | | Amide-I,PP-II |
